# Tempo and mode of diversification of the red devil spiders (Araneae: Dysderidae) of the Canary Islands

**DOI:** 10.1101/2024.07.09.601672

**Authors:** Adrià Bellvert, Laura J. Pollock, Antigoni Kaliontzopoulou, Miquel A. Arnedo

## Abstract

The study of adaptive radiations has shed light on our current understanding of evolution. However, previous studies examining the mode in which species diversified, how diversification rates varied, and how ecological specialization affected these processes have found few different results across different taxa and geographic and ecological systems, showing how complex this process is. To gain a more complete picture of how species evolve, additional model systems that encompass alternative ecological requirements are needed. Here, we present the results of a study aimed to unravel the diversification mode and evolutionary drivers of the spider genus *Dysdera*, the red devil spiders, endemic to the Canary Islands. These species exhibit remarkable phenotypic variability in their mouthparts, which has been related to different levels of specialization in the predation of isopods. We explored patterns of linage diversification and assessed the role of trophic specialization as a driver of species diversification. Additionally, we used climatic variables, occurrence data and morphological information to unravel the underlying mode of speciation by means of joint species distribution models and age-range correlation methods. Our results reveal that red devil spiders underwent an early burst of diversification, followed by a slowdown of diversification rates, which is a hallmark of adaptive radiation. We also found evidence that the trophic morphology shaped diversification, with specialist species exhibiting higher rates of diversification. Finally, our analyses suggest that speciation occurred mostly in allopatry, with subsequent secondary sympatry following range expansion.

## Introduction

Evolutionary radiations, or periods of accelerated species diversification, have been a prominent feature of life’s history on Earth (Linder & Bouchenak-Khelladi 2017). Among these, adaptive radiations – the diversification of ecologically and phenotypically distinct species from a common ancestor (Futuyma 1998; Schluter 2000) – have garnered significant attention due to their importance in shaping the planet’s biodiversity (Glor 2010; Harmon *et al*. 2010). Not surprisingly, the study of one of these radiations, the Darwin finches in the Galapagos Islands, was seminal for the development of the theory of evolution by natural selection (Darwin 1859). Indeed, the study of these events offers a unique opportunity to explore the mechanisms of species evolution and the complex relationships among extant species diversity (Givnish & Sytsma 1997; Losos & Mahler 2010). While adaptive radiations are observed in various environments, young and isolated geographic regions such as volcanic islands or glacial lakes offer ideal conditions for studying these phenomena (Schluter 2000) and have been instrumental in advancing our understanding of speciation and other evolutionary processes (Gillespie *et al*. 2020). As a result, many of the most well-known cases of adaptive radiation have occurred in isolated regions (Harmon *et al*. 2010; Patton *et al*. 2021; Cerca *et al*. 2023), including *Tethragnatha* spiders in the Hawaiian Islands (Gillespie 2004), cichlid fishes in the African lakes (Takahashi & Koblmüller 2011), the anoles lizards in the Caribbean (Losos 2009) and the aforementioned Darwin’s finches from the Galapagos (Lack 1947), to mention just a few. Hypotheses on what triggers these adaptive radiations have been mainly focused on ecological opportunity, i.e. the availability of new ecological resources not previously accessible (see Stroud & Losos 2016 for a review). Such ecological opportunities may emerge under different circumstances (Simpson 1953), but the colonization of oceanic islands, which are usually more species-depauperate, have provided the clearer examples (Stroud & Losos 2016; Gillespie *et al*. 2020). In this context, specific traits that allow linages to enter new “adaptive zones” (Simpson 1944) and promote ecological specialization (Heard & Hauser 1995), the so called “key innovations”, have also been a central element in the discussion of adaptive radiations, as they have been linked to increased species diversification rates (Simpson 1944; Mayr 1963; Losos & Mahler 2010; Wellborn & Langerhans 2015).

Despite the importance of adaptive radiations in shaping Earth’s biodiversity, our understanding of the detailed underlying mechanisms remains incomplete. While the evolution of ecologically and phenotypically distinct species from a common ancestor is the signature of adaptive radiations (Futuyma 1986), few other common processes have emerged from studies of these patterns. For instance, exploring associations between lineage and phenotypic diversification has produced mixed results when studying continental adaptive radiations, with asymmetric diversification between these two patterns (see Pincheira-Donoso *et al*. 2015), and the typical pattern of an early burst followed by a slowdown in linage diversification (Phillimore & Price 2008; Rabosky & Lovette 2008; Harmon *et al*. 2010), has lacked support in cases where secondary factors could have erased this typical signature (Slater *et al*. 2010). Such inconsistencies suggest that the dynamics of adaptive radiations are more complex than previously thought. Similarly, studies of the macroevolutionary consequences of ecological specialization on species diversification have also yielded disparate results. Historically seen as an evolutionary dead-end, specialist species, or lineages, have been hypothesized to exhibit lower diversification rates than generalists, evolving from them but not in reverse (Schluter 2000; Day *et al*. 2016). However, several studies have shown that both irreversibility and lower diversification rates assumed for specialist species are far from being a general pattern (Nosil 2002; Nosil & Mooers 2005).

The actual drivers of speciation are another potential source of conflict among different adaptive radiation processes. Historically, the widespread view was that speciation was mainly triggered by geographic isolation (Mayr 1963), that is, obliteration of gene flow coupled with time. However, with the rise of molecular phylogenetics and the use of more sophisticated comparative methods, that allows for quantitative detection of post-speciation range shifts (e.g. age-range correlation methods), it has become evident that the actual mechanisms underlying geographic patterns of species diversification could be more complex. Indeed, several studies have found evidence of sympatric speciation and thus of the potential involvement of natural or sexual selection in the actual speciation process (Bolnick & Fitzpatrick 2007). Comparative methods assessing the relationship between distribution overlapping and phylogenetic relatedness, are based on the assumption that, following the initial speciation stage, the geographical distribution of species becomes randomized due to their dispersal capabilities (Barraclough & Vogler 2000). In cases where a species clade undergoes allopatric diversification, the overlap between their ranges would initially be zero. However, over time, as their ranges randomly change, the overlap increases, resulting in a positive correlation between range overlap and older stages in the species’ phylogenetic relationships, relative to more recent cladogenetic events (Fitzpatrick & Turelli 2006). However, it is important to note that these correlations can be misleading when species clades exhibit complex patterns of geographical speciation (Fitzpatrick & Turelli 2006). For example, the randomizations in post-speciation range shifts may be compromised when species experience secondary contact after the original speciation process, as observed in numerous cases of island clade diversification (e.g. Thorpe *et al*. 2010; Macías-Hernández *et al*. 2013).

Given the intricate temporal, ecological and geographic variation in species diversification patterns and processes observed across different species groups, additional model systems that encompass alternative ecological requirements are needed to obtain a more complete understanding of how species emerge and evolve. The Canary Islands form a volcanic archipelago that includes seven major islands and several islets located 100 kilometres of the north-west of the African coast (Fig. 1a). The islands are geochronologically arranged, with the oldest ones, Lanzarote and Fuerteventura (15 My and 23 My respectively), lying at the easternmost side, and becoming progressively younger towards the western end (from east to west): Gran Canaria (subaerial age 15 My), Tenerife (12 My), La Gomera (11 My), La Palma (1.7 My) and El Hierro (1.1 My) (Van Den Bogaard 2013). These islands have been home to a remarkable diversification in the red devil spiders of the genus *Dysdera* (Fig. 1b), with approximately 60 endemic species recorded in the archipelago (Bellvert *et al*. in prep) out of a total of 300 currently described species (World Spider Catalog 2023). These species exhibit remarkable phenotypic variation in their cheliceral morphology—the spiderś mouth parts, which has been linked to different levels of trophic specialization (Řezáč *et al*. 2008, 2021; Toft & Macías-Hernández 2017, 2021). Different cheliceral morphologies evolved independently several times during the diversification of the group in the islands ( Bellvert *et al*. 2023). Because of the single colonization event inferred for these species (but see Adrián-Serrano *et al*. 2020) and the eco-phenotypic differences observed among relatives, this genus has been suggested to be a case of adaptive radiation in the islands (Bellvert *et al*. 2023), but this hypothesis has never been formally tested and the exact evolutionary drivers of this diversification remain poorly understood.

**Figure 1.**
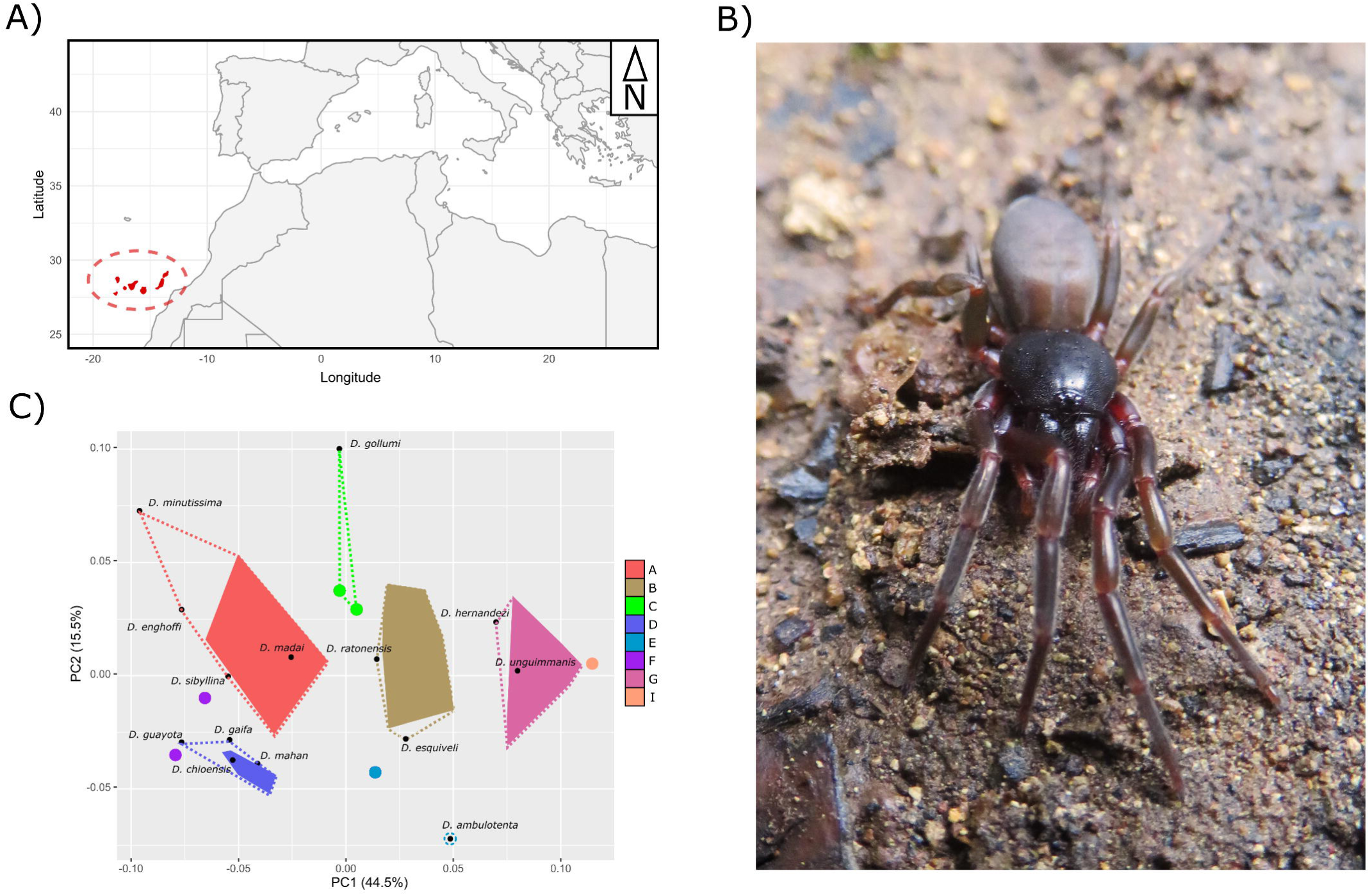
a) Map of the Canary Islands. b) *Dysdera verneaui* from the island of Tenerife (photo credit: Marc Domènech). C) Cheliceral morphospace obtained with the two first principal components from the morphometric data used for this study from Bellvert et al. (2023), with densely-colored convex polygons depicting species of the same ecomorph group. Black dots represent the species not included in the previous study but with morphometric data. Colored dots represent species already previously linked to cheliceral shape but without enough species to build a polygon. Doted lines represent the change in the cheliceral type polygon space once the species have been assigned to the preexisting cheliceral types by means of the LDA performed in the present study.

In the present study, we used phylogenetic comparative methods and joint species distribution modelling to elucidate the mechanisms underlying the diversification of *Dysdera* spiders in the Canary Islands and examine how it has been influenced by trophic specialization. Our specific objectives were: 1) to test if the temporal pattern of lineage diversification of the group aligns with that expected under a scenario of adaptive radiation, i.e. an early rapid diversification of linages with a subsequent deceleration (Glor, 2010); 2) to explore whether trophic specialization played a role in this diversification; and 3) to investigate whether allopatry followed by secondary overlapping drove the diversification of these species, similarly to other cases of adaptive radiation in oceanic islands. We conducted a comprehensive study by integrating the fully resolved phylogeny of the group with phenotypic data, occurrence records, and climatic variables. Our investigation aimed to understand the mode and tempo of the genus’ evolution in the archipelago, as well as to identify the factors that potentially played a role in its remarkable diversification.

## Methods

### Specimens and data collection

All specimens used came from field campaigns conducted by the authors and other colleagues and are stored at the Centre de Recursos de Biodiversitat Animal of the Universitat de Barcelona (CRBA) and Departamento de Zoología de la Universidad de La Laguna, Tenerife, Canary Islands collection (DZUL). Individuals were captured by active searching under rocks, logs and tree barks. The captured specimens were preserved in 95% EtOH and stored at -20°C at the former institutions. All specimens were collected following institutional and governmental regulations and the permits for all species captured were granted by the local authorities of each island or by the governing body of each natural reserve. Phenotypic data for the present study came from Bellvert *et al*. (2023). We used the statistical environment R (R Core Team 2023) to conduct all subsequent analyses.

### Time calibrated phylogeny

We inferred a time calibrated phylogenetic tree of Canarian endemic species and selected outgroups using mitogenomic data from Bellvert *et al*. (2023). We adjusted the taxon sampling and calibration points. We included only Dysderidae representatives, with the outgroup species *Opopaea cornuta* Yin & Wang, 1984 (Oonopidae) used to root the tree. We used a combination of 2 fossil calibrations and 5 biogeographic events to constrain nodes in the calibration information. The Baltic amber fossil *Dasumiana emicans* Wunderlich, 2004, has been classified as a member of the Harpacteinae (Bellvert *et al*. 2023). This information was used to establish a lognormal prior distribution for the Harpacteinae crown node, with an offset of 43 million years ago (Mya), which aligns with the earliest age of the Baltic amber (Magalhaes *et al*. 2020). Additionally, a hyperprior was set for the mean of the lognormal distribution, with a uniform distribution ranging between 0.01–40 Mya. The fossil *Burmorchestina acuminata*, identified as a member of the Onopidae subfamily Orchestininae, from Burmese amber was used to constrain the tree root. It was assigned a uniform prior ranging from 98.17 Mya, the presumed age of the Burmese amber and 164 Mya, the earliest evidence of a Synespermiata species, the major clade to which Dysderidae family belongs (Magalhaes *et al*. 2020). The biogeographic information included the Hercynian split of the Iberian plate into present-day major western Mediterranean islands, dated at 33–25 Mya (Rosenbaum *et al*. 2002; Schettino & Turco 2006). The information was used to set a normal distribution prior on the node splitting the Iberian and the island species of the genus *Parachtes* Alicata, 1964, with mean 29 Mya and standard deviation 2.5 (Bidegaray-Batista & Arnedo 2011). The remaining biogeographic constraints were provided by the estimated emergence of the younger Canarian Island in species with sister populations on more than one island. Specifically, the species *Dysdera gomerensis* Strand and *D. silvatica* Schmidt have sister lineages in La Gomera and El Hierro, respectively. The earliest subaerial age of El Hierro, the younger island, has been estimated at 1.2 My (Carracedo & Day 2002). We defined a uniform prior ranging from 0.1–1.5 Mya to the two nodes splitting the former species pairs, to account for both earlier splits and later colonization of the youngest island. Similarly, the species *D. calderensis* Wunderlich, 1987and *D. silvatica* have sister lineages in La Gomera and La Palma. The earliest subaerial age of La Palma has been estimated at 2 My (Carracedo & Day 2002), and we defined a uniform prior ranging 0.1–2.5 Mya to both nodes.

We inferred the time-stamped calibrated phylogeny under a Bayesian uncorrelated relaxed molecular clock approach as implemented in BEAST v2.7.6 (Bouckaert *et al*. 2019). The concatenated data matrix was partitioned by gene, and the best evolutionary model for each gene partition was selected with PartitionFinder 2 (Lanfear *et al*. 2017). Individual log-normal clocks were defined for each gene, and the tree prior was set to the Birth-Death model.

We run three independent chains under selected priors for 100 million generations, sampling every 10,000 generations. Convergence among runs and correct mixing of the chains was monitored with TRACER v.1.7.1 (http://tree.bio.ed.ac.uk/software/tracer/). The burn-in was removed (10%) and the runs were combined with the help of the BEAST accompanying programs LOGCOMBINER and TREEANNOTATOR.

### Species linage diversification

To investigate the diversification mode in the *Dysdera* spiders from the Canary Islands, we used the time calibrated phylogeny, following removal of non Canarian species and the endemic *D. lancerotensis*, as it has been shown to correspond to an an independent colonization not directly related with the rest of the Canarian species (Adrián-Serrano *et al*. 2020). To obtain a first view on how diversification proceeded across the history of the group, we extracted the branching times with the function *branching.times* from the R package “ape” (Paradis & Schliep 2019) and fitted different variable-rate and constant-rate models to this data using the function *fitdAICrc* from the R package “laser” (Rabosky & Schliep 2013) considering 100 shift points and 6 different models: pure birth, birth-death, an exponential and a logistic variant of density-dependent and speciation rate models, pure birth with a single rate shift and pure birth with two rate shifts. To visualize the number of lineages through time and obtain a summary of diversification rates we used a linage-through-time plot produced with the function *ltt* from the R package “phytools” (Revell 2012). To consider phylogenetic uncertainty we repeated these analyses over 1000 trees obtained from the posterior distribution from the study of Bellvert *et al*. (2023). Previous work has delimited, based on geometric morphometric methods, several cheliceral morphotypes that have evolved independently several times during the diversification of these species in the archipelago (Bellvert *et al*. 2023), and which are hereby hypothesized to have potentially driven species diversification, specifically through their effect on the species trophic specialization, where certain cheliceral morphologies are associated with generalist predators, while other morphologies exhibit varying degrees of specialization in preying on isopods. However, not all the *Dysdera* diversity from the Canary Islands was represented in the aforementioned study, which only included 40 out of a total of 57 currently recognized species (Bellvert *et al*. 2023). Such undersampling could lead to a bias when analyzing evolutionary rates (Pybus & Harvey 2000) that could compromise the final results of the study. Some of the unrepresented species were due to lack of any specimen to obtain morphometric data; and had to be excluded from the study. However, other specimens where previously ignored because of a lack of intraspecific variation or because they belonged to other ecological regimes (see Bellvert *et al*. 2023). To mitigate the effect of the undersampling on the diversification analyses conducted here, we used a Linear Discriminant Analysis (LDA) to classify the species with unknown cheliceral affiliation but with some morphometric data into the already established cheliceral morphotypes. We used the function *prep.lda* from the R package “RRPP” (Collyer & Adams 2018, 2021) to prepare the geometric morphometric data used in Bellvert *et al*. (2023) study and the function *lda* from the R package “MASS” (Venables & Ripley 2002) to run the discriminant analysis. For subsequent analyses, we pruned the complete phylogeny of the *Dysdera* spiders from the Canary Islands with the species with cheliceral type information from Bellvert *et al*. (2023) and the ones that we could recover from the discriminant analyses. We used the function *drop.tip* from the R package “ape” (Paradis & Schliep 2019) to remove all specimens without cheliceral type information from the phylogeny.

### State dependent diversification models

To test if tropic specialization as represented by different cheliceral morphotypes increased or decreased the rate at which *Dysdera* species diversified, we fitted a BiSSE model where diversification and extinction rates varied freely between two distinct states of trophic strategy using the function *make.bisse* from the R package “diversitree” (Fitzjohn 2012). Binary states were obtained from Bellvert *et al*. (2023) which link the different cheliceral morphologies to generalist or specialist trophic strategies. We compared this model to a null model with equal diversification and extinction rates by extracting the loglikelihood of each model through the function *find.mle* and comparing both with the function *anova*. Furthermore, to test and visualize differences in diversification rates between trophic strategies, we used Bayesian inference to calculate the posterior distribution of our parameters with the function *mcmc* from the R package “coda” (Plummer *et al*. 2006).

However, it could be the case that cheliceral morphotype by itself, capturing the evolutionary influences of other pressures other than or in addition to trophic strategy, could have affected diversification rates. In order to test if the evolution of different cheliceral morphotypes, rather than their trophic strategy, have an impact on the diversification rates in the *Dysdera* species, an optimal approach would be to use a multi-state speciation and extinction model like MuSSE (Fitzjohn 2012). Unfortunately, in our data some cheliceral morphologies were only represented by one or two species, and the resulting MuSSE model would probably not be trustworthy. Instead, to investigate which cheliceral types, or combination of some of them, could have had a bigger influence on the *Dysdera* diversification, we ran independent BiSSE models for each possible binary combination among all 8 cheliceral morphotypes. From each of these models, we calculated the percentage of times that each cheliceral type appeared in pairwise cheliceral combinations that were inferred to exhibit significantly different diversification rates when compared to a null model with equal rates between states. We represented the percentage of times that each cheliceral morphotype appeared in significant pairwise combinations with significantly different diversification rates using a spider chart with the function *radarchart* from the R package “fmsb” (Nakazawa 2023). Finally, to examine which cheliceral combination maximized change in diversification rates, we ran a BiSSE model delimiting the binary state with the cheliceral types that showed a higher percentage of occurrence in the previous combinations. We compared a model with different diversification and extinction rates with a null model with equal rates, with the binary state representing each of the three cheliceral types with the highest percentages of occurrence on the one hand and the rest of types on the other. Next, we also examined a model with the binary state defined as the five cheliceral types with the highest percentages and the two cheliceral types that have not appeared a single time in the combination of cheliceral morphologies that significantly increase diversification rates (see results).

Note that a trait-state-dependent model with different diversification rates that performs better than a null one, does not necessarily mean that the examined trait is the main driver of the diversification of the group, as an unmeasured factor could have a stronger impact on the species’ diversification (Beaulieu & O’Meara 2016). To account for this possibility, for all previous models with the binary state delimited based on the cheliceral percentages in which the different-rates model performed better than the null, we fit three additional “hidden rate” models (Beaulieu & O’Meara 2016): one in which there is an influence on diversification rates of an unaccounted trait but not of the cheliceral type (CID-2), a four-level hidden rate model (CID-4), which will allow to test for more complex diversification processes (Revell & Harmon 2022), and a full model were both the hidden state and our cheliceral trait have an effect on the Canarian *Dysdera* spider diversification. We used the *hisse* function from the R package “hisse” (Beaulieu & O’Meara 2016) to fit these models. We calculated the AIC of each model with the *AIC* function to test which performed best.

### Time dependent diversification models

We tested if diversification rates varied through time in the radiation of the *Dysdera* spiders from the Canary Islands. We used the function *make.bd.t* from the R package “diversitree” (Fitzjohn 2012) to fit a model were diversification rates were an arbitrary function of time. We compared the time-dependent model to a null with a constant diversification rate fitting our data with the function *make.bd*. We obtained the maximum likelihood of each model with the function *find.mle* and compared between them with the *anova* function. We also used the *mcmc* function to run a Bayesian MCMC analysis for our time dependent model and plot the posterior distribution through time of speciation (λ) to see the general tendency that this parameter has.

### Species overlap and age-range correlation

Finally, to unravel the main speciation pattern of these species in the islands—i.e. allopatric vs sympatric—we analyzed how the overlap between species distribution ranges changed through time. We used Age-range correlation methods (ARC), which seek to establish statistical relationships between species-pairs geographic overlap and the time since their divergence (Fitzpatrick & Turelli 2006). For the pairwise measurements of the niche overlap between species, we used joint species distribution modelling (jSDM) to obtain the residual patterns of occurence between species, as they capture more complex interactions than other co-ocurrence metrics could show (Pollock *et al*. 2014). Those residuals are features that influence occurrence between species that are not explained by environmental variables. We decided to use these residuals, instead of the species shared environmental responses, because they explain better the difference between allopatric and sympatric species pairs (see results). We used all the species for which we had sufficient morphometric data from Bellvert *et al*. (2023) (N = 40), and we considered the first two principal components of each species as the trait value for the model to also test for competitive exclusion in the biotic interactions captured by the residual correlations. For the species’ occurrence, we built a presence matrix for each species for each locality were *Dysdera* spiders have been collected in the Canary Islands during the last 50 years. Finally, for the model explanatory variables, we selected the 3 different climatic predictors from Patiño *et al*. (2023) that we hypothesize may have a greater impact on the species’ potential distribution, namely the highest temperature of any month, to include the potential effect of aridity; the accumulated precipitation over 1 year, to take into account the effect of the more humid areas; and elevation, which has an influence on temperature but also on the vegetation present in the island. We ran the model with the function *jSDM_binomial_probit* from the “jSDM” R package (Clément & Vieilledent 2022), with 10000 iterations with a burn-in of 5000 and a thinning rate of 20. We extracted the latent variables of the model and the correlation of the response matrix with the *get_residual_cor* and *get_enviro_cor* functions respectively. For the ARC analysis we used with the function *age.range.correlation* from the R package “phyloclim” (Heibl & Calange 2018). An observed negative slope to more distant relationships would point to a sympatric diversification, while a positive one would be indicative of an allopatric diversification (Fitzpatrick & Turelli 2006). We ran the model with 999 iterations to test for the statistical significance.

## Results

### Best macro-evolutionary candidate model

The gamma statistic exhibited a negative value (-5.35 ±0.18), being significantly smaller than the expected under a birth death process which would have a normal distribution around 0 (Fig. 2a), which suggests rejection of the hypothesis of constant rate in the diversification of the group in favor of decreasing diversification through time (Pybus & Harvey 2000). Accordingly, the LTT plot showed a clear hump-shaped increase in the number of lineages with a posterior stabilization (Fig. 2b). Finally, when contrasting different candidate diversification models, the density-dependent speciation model (DDL) showed the best performance with an AIC of -37.76±5.06, followed by the pure-birth model with two rate shifts (yule 3 rates, AIC = -33.73±5.41), the pure-birth with one shift (yule 2 rates, AIC = - 33.65±5.79), the exponential variant of the density-dependent speciation model (DDX, AIC = - 18.89±4.50), the pure-birth model (AIC = -6.62±4.59) and the birth-death model (AIC = -4.62±4.59).

**Figure 2.**
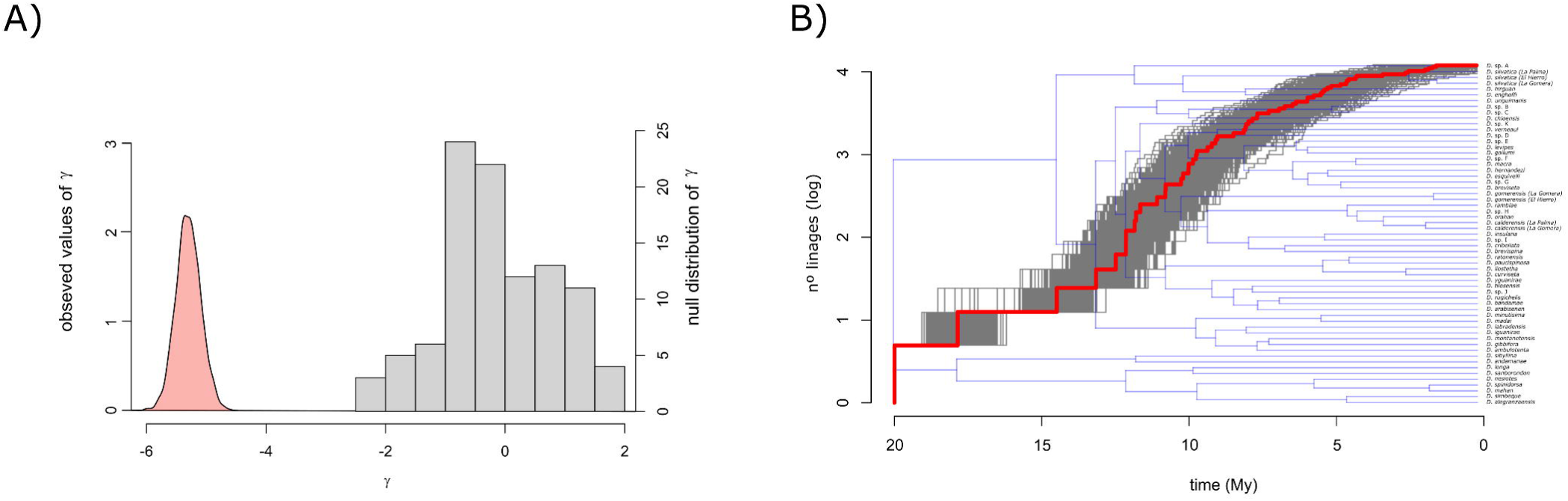
a) observed γ values for 1000 phylogenetic trees obtained from the posterior distribution (red), compared to the γ values from a null distribution under a pure birth model (grey histogram). b) LTT plot for the *Dysdera* species from the Canary Islands. The red line represents the LTT made with the consensus tree, grey lines are the LTTs from the 1000 phylogenetic trees from the posterior distribution used to take phylogenetic uncertainty into account.

### BiSSE state-dependent models

The results from the discriminant analysis and the cheliceral type adscription can be found in the supplementary material and in Table S1. The BiSSE model with trophic strategy as the binary state (specialist vs generalist) with free diversification rates (fr), performed poorly compared to the null model with equal rates between states (null), without significant differences between them (AIC_fr_ = 364.05, AIC_null_ = 362.81, p-value = 0.25). However, the Bayesian MCMC analysis disagreed with the previous one and showed that the diversification rate of specialist species is higher than that of generalist ones (p-value = 0.046). When examining all pairwise combinations of binary states among cheliceral types, we found that for 17 out of 254 possible pairwise combinations the model with free diversification rates performed better as compared to the null model with equal rates between states (Table S2). From all these 17 different cheliceral combinations, type B was among the cheliceral morphologies that exhibited a higher diversification rate in 89% of the cases, type F in 55% of the cases, type G in 50% of the cases, type C and E in a 39%, type I in 33% of the cases, while types A and D were never part of the higher diversification rate sets. When creating groups of several cheliceral types to generate the binary state trait for rate comparisons depending on those percentages, we found that, when comparing the cheliceral morphotypes that appeared at least 50% of the times in the cheliceral combinations that showed an increase in diversification rates against the other ones (i.e. B, F and G vs C, E, I, A and D), the BiSSE model with free transition rates exhibited a better performance than the null (AIC_fr_ = 366.23 and AIC_null_ = 368.23, p-value = 0.049), with the Bayesian MCMC analyses showing that state_BFG_ have higher diversification rates compared to state_ICFAD_ (Fig. 3a, p-value = 0.014). However, when combining the cheliceral morphologies that appeared less than 50% of the times in the previous combinations (state_BFGCE_), against the cheliceral morphologies that does not appeared a single time in the state with an increase in diversification rates (state_AD_), the model with free transition rates does not performed better than the null model (AIC_fr_ = 373.82 and AIC_null_ = 371.28, p-value = 0.48), with the Bayesian MCMC analyses not showing differences in the diversification rates between state_BFGCE_ and state_AD_ (Fig. 3b, p-value = 0.066).

**Figure 3.**
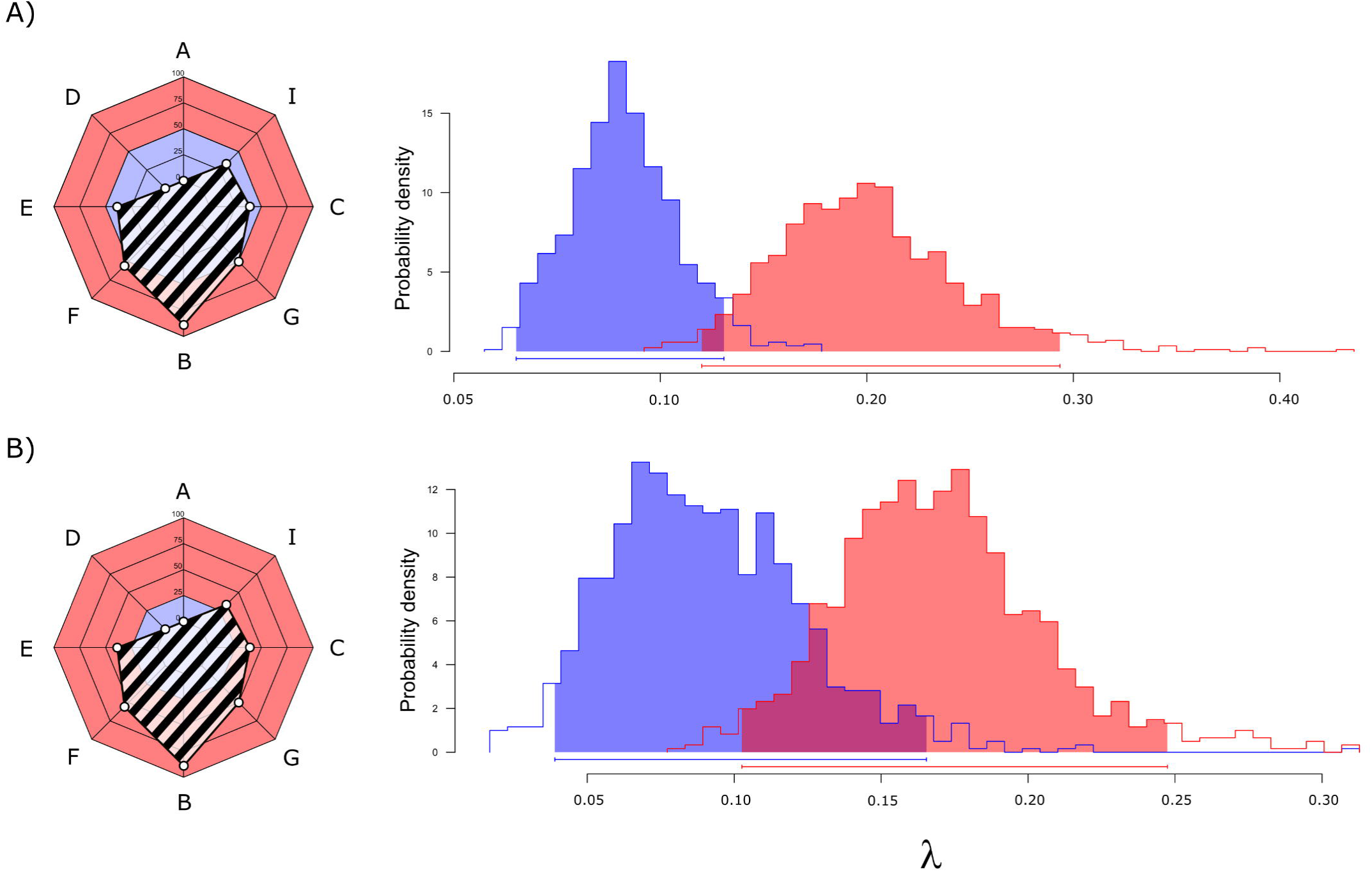
Diversification rates obtained with the BiSSE test for binary states a) state_BFG_ (red) and state_ICFAD_ (blue); b) state_BFGICE_ (red) and state_AD_ (blue). The spider chart shows the binary state delimitation with the percentage of occurrence for each state in the binary states’ combinations (black striped).

### Hidden state results

When testing for a hidden state character responsible for differences in diversification rates among Canarian *Dysdera* species, we found that models accounting for one or more hidden characters, or a combination of a hidden character and the cheliceral morphotypes, performed poorly compared to the model focusing solely on differences in diversification rates based on cheliceral morphology (state_BFG_ vs state_CEIAD_, BISSE AIC = 366.22, CID AIC = 368.23, HISSE CID-2 AIC = 373.56, HISSE CID-4 AIC = 377.39, HISSE+BISSE AIC = 391.99).

### Effect of time on species diversification

When investigating temporal variation in diversification rates, we found that the time-dependent model performed better than the null with no variation in diversification rates (AIC = -21.23 and AIC = 22.12 respectively, p-value << 0.01). When sampling the posterior distribution using Bayesian inference, we observed a slight decrease in the diversification rate of the group through time (Fig. 4, lambda.a = 0.0069 ± 0.0073).

**Figure 4.**
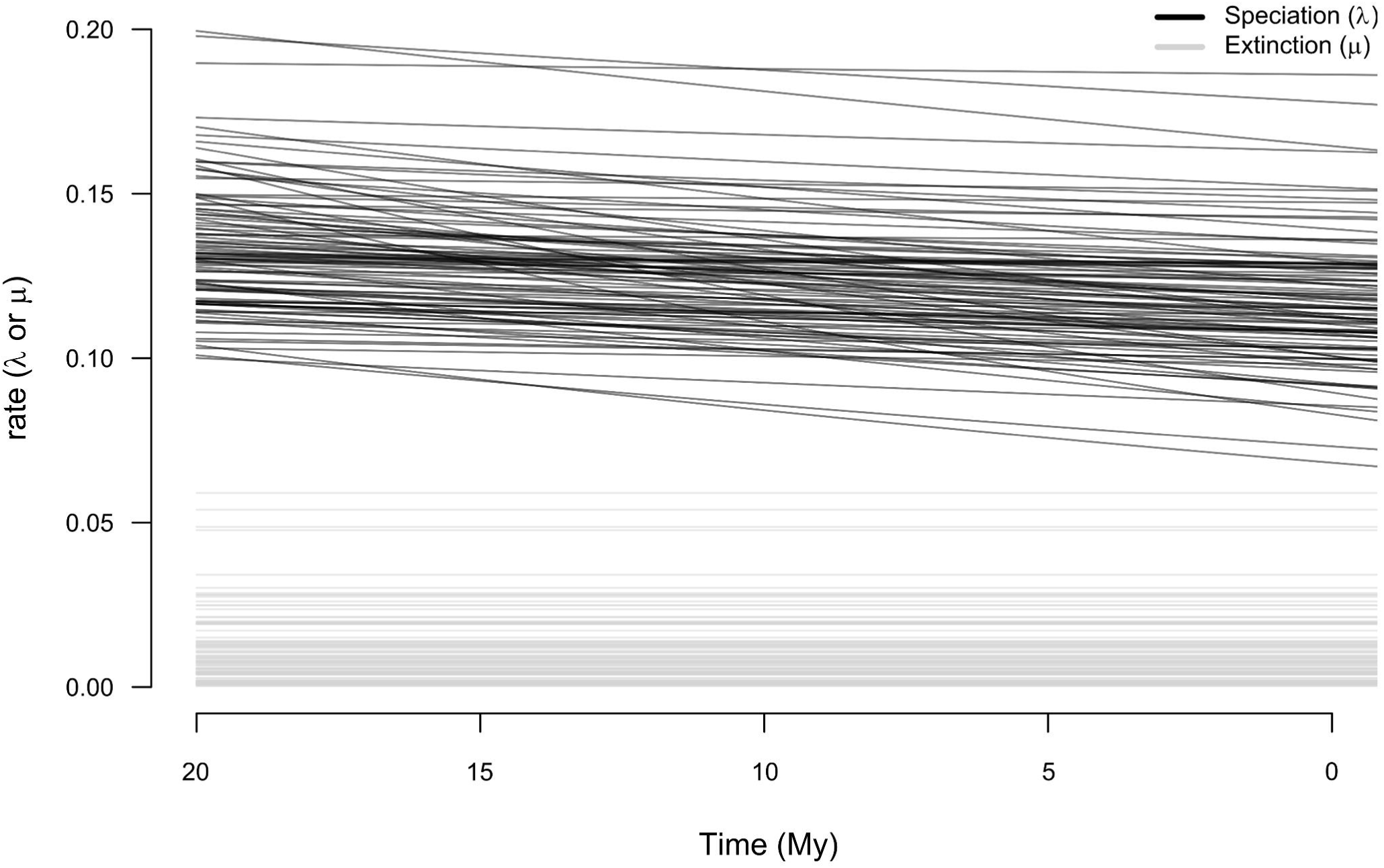
Posterior distribution of the time-dependent speciation and extinction rates obtained with the variable-speciation model.

### Age-range correlation

jSDM showed varying correlations between different species pairs, both in terms of their correlation with the environmental variables and their residual correlations. However, upon examining species pairs that occur on the same island, we found that they generally exhibited positive correlations with both residuals and environmental factors. Conversely, for species pairs that occur on different islands, the correlation with the residuals tends to be neutral but with a bigger disparity in the environmental correlations, with both positive and negative values (Fig. 5a). We opted to utilize residual correlations for conducting the ARC analyses, as they provide better discrimination between species pairs with distributions on the same or different islands. It is important to note that having a distribution on the same island does not necessarily imply a sympatric distribution, as the species may occupy different regions within that island. However, the residual correlations offer the clearest representation of what an allopatric distribution could be (i.e., two species on separate islands), and because of that we believe they will contribute more significantly to the ongoing analysis.

**Figure 5.**
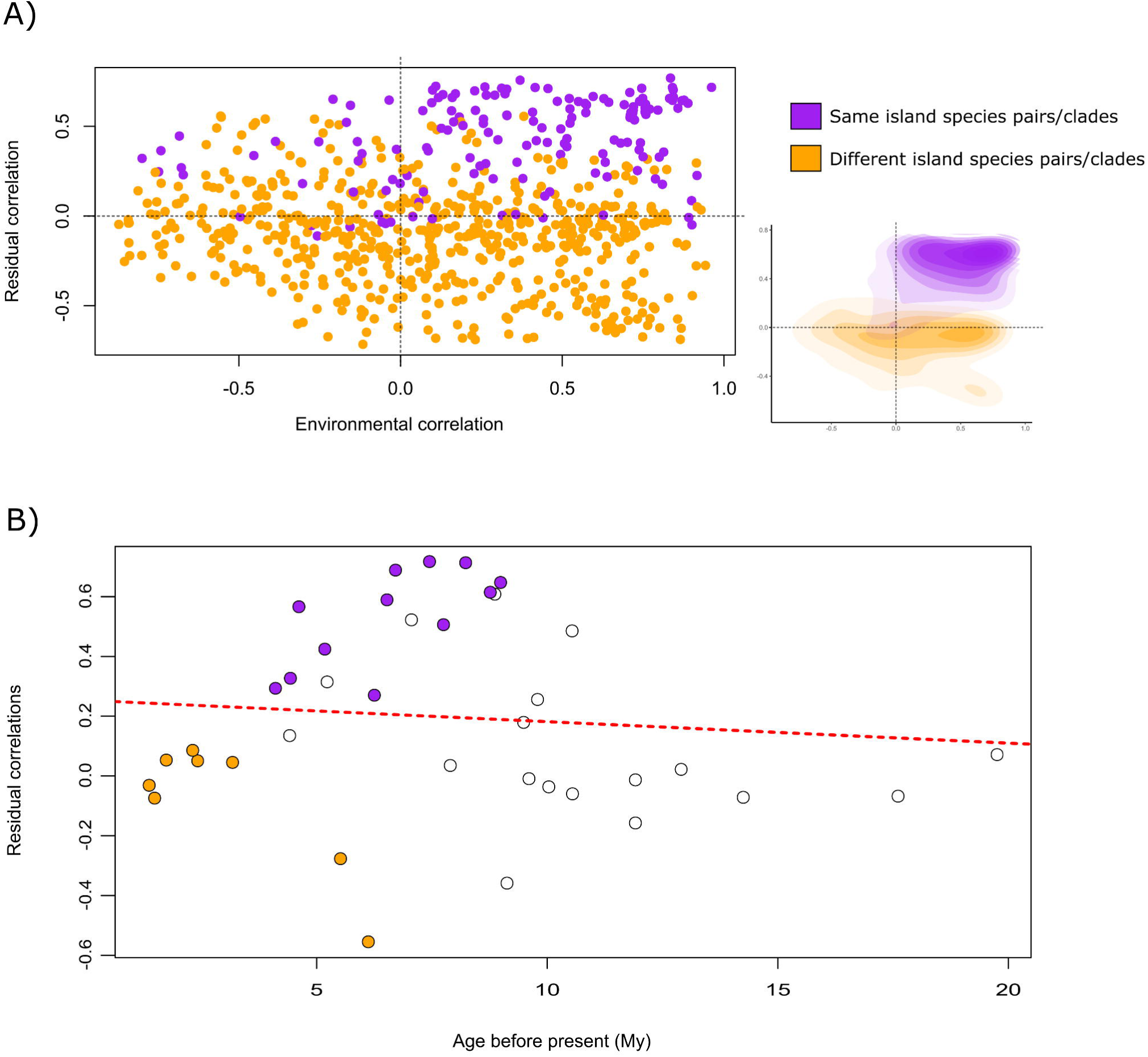
a) Plot with residual correlations and environmental correlations between all *Dysdera* species pairs in the Canary Islands. Purple dots represent species pairs that inhabit the same island, and orange dots species pairs that have a distribution on different islands. Bottom right corner represents the density concentration of sympatric and allopatric species pairs in the residual and the environmental correlations, showing bigger differences between their densities in the residual correlations. b) Age-range correlation of node ages and species residual correlations obtained from the jSDM. Doted red line represent the linear model of mean residual correlation versus node age. Dots represent nodes in the phylogeny: orange dots are sister species pairs that occur on different islands; purple dots are species pairs or clades that occur on the same island; white dots are nodes that cannot be linked because the species relations are too complex to characterize the node as sympatric or allopatric.

The ARC analysis revealed a decrease in the overlap since species divergence (intercept = 0.253; slope = -0.007), indicative of a sympatric speciation with a subsequent loss of contact between species. However, this result was not statistically significant with a p-value of 0.366, showing no significant association between range overlap and time since species divergence. It should be noted, however, that these values were obtained by fitting the results in a linear model, but our ARC plot clearly exhibited a hump-shaped pattern, with low overlap values in the early splits, a subsequent increase in overlapping followed by a decrease toward the present (Fig. 5b).

## Discussion

In this study, we have focused in understanding how species diversification varies across time and space, and how it is influenced by ecophenotypic traits and trophic specialization. Moreover, we have provided evidence that the diversification of *Dysdera* spiders in the Canary Islands, where previous studies have highlighted eco-phenotypic variation (Bellvert *et al*. 2023), constitute a case of adaptive radiation. Evidence derived from modelling speciation and extinction lends support to the hypothesis that trophic specialization and associated morphological evolution shaped the diversification of these species in this archipelago, highlighting the importance of ecological innovation in driving species’ diversification. Finally, we have found evidence that the speciation of the *Dysdera* species in the Canary Islands had been mainly driving by geographic isolation with a posterior secondary contact.

### The diversification mode of Canarian *Dysdera* spiders

An early increase in species diversification, followed by a decrease in net speciation rates towards the present is the trademark of an adaptive radiation process (Phillimore & Price 2008; Glor 2010; Losos & Mahler 2010). Declining diversification rates through time are generally interpreted as the results of early occupation of vacant niches until their saturation, which leads to the slowdown of cladogenetic events (Schluter 2000; Gavrilets & Vose 2005). Such a pattern has been usually described by means of linage-through-time plots (Nee *et al*. 1992; Pybus & Harvey 2000), with density-dependent models describing this asymptotic accumulation of lineages through time (eg. Pincheira-Donoso *et al*. 2015; Linder & Bouchenak-Khelladi 2017). The early increase in number of linages and the corresponding negative γ values revealed for the *Dysdera* species from the Canary Islands, points towards a rapid filling of ecological niches and a deceleration of speciation rates over time, as also seen with the time-dependent model of diversification. Interestingly, the Canary Islands may have also presented additional episodes of ecological opportunity as new islands emerged over time, which can be reflected in diversification pulses in the LTT plot (Pincheira-Donoso *et al*. 2015). Such pulses can be identified in Fig. 2b. Multi-rate variants of the pure birth model lend further support to this observation, as they exhibited overlapping AIC values when accounting for the phylogenetic uncertainty, however, their mean values has been slightly higher than the density-dependent model. Obviously, undiscovered species could compromise these results, as incompleteness in the taxon sampling could mirror a similar pattern as the density-dependent speciation (Rabosky & Lovette 2008). However, given the solid taxonomic ground established by a series of modern taxonomic treatments of the group (eg. Arnedo & Ribera 1999a, 1999b, 1999c; Arnedo *et al*. 2000; Macías-Hernández *et al*. 2010; Bellvert *et al*. 2023), we doubt that the number of undiscovered species may actually affect our main conclusions.

### Evolutionary consequences of ecological specialization

The ecological and evolutionary consequences of species specialization have been examined repeatedly over the years. Current consensus would claim that the evolution of traits that reduce the breadth of species trophic or ecological niches negatively affects their diversification rates (Vamosi *et al*. 2014), resulting into the so-called evolutionary dead-ends (Cope 1896). However, this cannot be taken as a general rule, as many studies have found mixed support for it (Day *et al*. 2016), or directly contradicted the link between specialization and evolutionary dead-ends (Zenil-Ferguson *et al*. 2022). Our results reveal that, globally, specialist and generalist *Dysdera* species exhibit similar diversification rates. Nevertheless, pairwise rate comparisons of different cheliceral morphotypes and their combination into rate-coherent groups identified specific sets of ecomorphotypes that exhibit significantly higher diversification rates (Fig. 3a-b). Interestingly, although trophic specialization does not seem to be directly associated to an increase in diversification rates in our studied species, we found a significant increase when combining cheliceral morphologies that have been previously linked to specialist trophic strategies (Bellvert *et al*. 2023). The group including cheliceral morphotypes B, G and F seem to exhibit elevated species diversification rates, while other cheliceral morphologies, some also related to a specialist diet (C and I), do not show this acceleration. However, the rare evolution of cheliceral type F across the Canarian *Dysdera* diversification, where this specific morphotype has only appeared in two independent occasions, makes its contribution to the inferred diversification dynamics questionable. Indeed, the increase in diversification rates attributed to cheliceral type F could be an artifact due to the close relatedness of these species to those with cheliceral morphologies B and G. In fact, in continental *Dysdera* this cheliceral type (F) has also evolved multiple times across different species groups and geographic regions, presumably independently. Here, this morphotype is generally carried by single species with no close relatives exhibiting the same feature, which also reinforces the idea that most likely this cheliceral morphology is not a key innovation responsible for the increase of diversification rates in *Dysdera* species.

In a previous study, we found transitions from generalist cheliceral morphotypes to specialist ones to be irreversible in Canarian *Dysdera* (Bellvert *et al*. 2023), supporting the hypothesis that prey specialization is an evolutionary dead-end (Day *et al*. 2016). However, the high diversification rates found with the specialist cheliceral morphologies B and G, do not support the idea of an evolutionary dead-end, suggesting instead that trophic specialization in *Dysdera* species may constitute a key innovation rather than a limiting factor in their diversification. On the other hand, the cheliceral morphologies C and I, and most likely also the cheliceral type F, would better fit into the definition of a dead-end, being an irreversible character that has evolved from a generalist state and that is also accompanied by a decrease in species diversification rates (Fig. 3c). Put together, these observations suggest that the evolutionary dead-ends in *Dysdera* spiders would not be so dependent on trophic specialization *per se*, but rather related to specific morphotypes.

A point of relevance here is that, for complex diversification processes, BiSSE models are known to frequently correlate neutral traits with higher diversification rates (Rabosky & Goldberg 2015; Beaulieu & O’Meara 2016). In spiders, it has already been observed that, although ecologically important, some traits may not be the primary driving force behind species diversification (Fernández *et al*. 2018). However, our analyses provide strong evidence on the relevance of morphotype differentiation for species diversification: when we used HiSSE to test whether differences in diversification rates might be explained by other unaccounted factors, we found that BiSSE models performed better than the CID-2 or CID-4 models. This indicates that cheliceral morphology indeed plays a strong role in the evolutionary history of these species.

### Secondary contact following allopatric speciation

The controversy of how generalized allopatric mode of speciation are compared to sympatric ones, especially in the context of adaptive radiations, is still a hot topic nowadays (see Bolnick & Fitzpatrick 2007). One of the major concerns about deciphering speciation with extant sister taxa has been the range shifts experienced by species that could blur present day patterns (Losos & Glor 2003), which were confronted by methods that account for post-speciation range shifts like ARC methods. The ARC analysis performed with the *Dysdera* spider species from the Canary Islands showed a non-significant negative correlation, related to a sympatric mode of speciation. However, the relation between age and the residual correlations has showed to be non-linear (Fig. 5b). Previous skepticism had already been raised in the past about these methods when analyzing complex patterns of geographical modes of speciation (Fitzpatrick & Turelli 2006). Sympatric patterns are generally explained no as the result of selection driven speciation, but as secondary contact following population expansion once the overlapping linages fully diverged genetically (Hudson *et al*. 2011). Our results of the ARC analysis would better fit under an allopatric speciation followed by a secondary sympatry, making the correlation presented in this test misleading of a more complex pattern that the *Dysdera* species may have been involved. The posterior decline in the species overlap observed could be a response to the increase of extinction rates expected over time under the general dynamic model of oceanic island biogeography (Borregaard *et al*. 2017). This allopatric speciation would suit into the expected in an archipelago, where intra-island geographical isolation could be less likely for these species than isolation by colonizing two separate islands (Mittelbach & Schemske 2015). This would match with the observed in our ARC analysis, where cladogenetic events within species pairs in different islands are more recent than the ones with species in the same island (Fig. 5b). For this reason, we argue that the colonization of different islands has been the main, or one of the main, speciation force(s) during the diversification of the *Dysdera* species in the Canary Islands

## Conclusions

The diversification of the *Dysdera* species in the Canary Islands has been previously suggested to be a case of adaptive radiation. Our results unambiguously support this claim by recovering an early burst of species diversification with a posterior slowdown in their speciation, a pattern usually interpreted as the stamp of an adaptive radiation process. The integration of available morphometric data and SSE models, provide evidence that the different cheliceral morphologies exhibited by Canarian *Dysdera* played a role in shaping diversification dynamics in this group, probably by increasing rates as a result of multiple instances of trophic specialization. The use of jSDM approaches provided more refined information on species associations than the use of climatic variables alone, and when used with ARC methods enabled us to propose that inter-island allopatric speciation is the general pattern of diversification among these species with sympatry being the results of secondary contact between them.

## Acknowledgments

A.B. was funded by an individual PhD grant BES-2017-080538 from the Ministerio de Economía, Industria y Competitividad of the Spanish government. A.K. is supported by a Ramón y Cajal research grant co-funded by the Spanish State Research Agency and the European Social Fund (RYC2019-026688-I/AEI/10.13039/501100011033). This study was supported by project grants CGL2012-36863 and CGL2016-80651-P from the Spanish Ministry of Economy and Competitivity and 2017SGR83 from the Catalan Government (M.A.).

