## supplementary material for "Tempo and mode of diversification of the red devil spiders (Araneae: Dysderidae) of the Canary Islands"

Linear discriminant analysis

From the 14 species for which we had morphometric data, but previous studies had no information about the cheliceral group, the LDA analysis assigned 13 to an already preestablished morphotype (posterior probabilities are summarized in Table S1 of supplementary material). The only species that could not be clearly classified to any group was *D. madai*. However, this species is morphologically very similar to other species of group A, as showed by its closeness in the morphospace represented by the first two principal components (Fig 1C). It is true that the differences with this species could be explained by other components, however the first two components explains 60% of the explained variability, and the posterior probabilities of the LDA showed similar values between the cheliceral group A and the unknown group (Table S1). For this reason, and to avoid over-splitting the number of different cheliceral morphotypes present in the Canary Islands, we considered this species to belong to the cheliceral morphotype A.
