## Supplementary material for "Tempo and mode of diversification of the red devil spiders (Araneae: Dysderidae) of the Canary Islands": Table S1

Table S1: LDA posterior probabilities for each unknown species cheliceral morphology to be adscrived to one of the already known types.

| Species | Group A | Group B | Group C | Group D | Group E | Group F | Group G | Group I | Unknown |
| --- | --- | --- | --- | --- | --- | --- | --- | --- | --- |
| *D. ambulotenta* | 0.00530 | 0.14003 | 0.00093 | 0.05322 | 0.41971 | 0.00058 | 0.12063 | 0.00335 | 0.04007 |
| *D. chioensis* | 0.16418 | 0.00510 | 0.00149 | 0.46749 | 0.01348 | 0.14922 | 0.00001 | 0.00000 | 0.19902 |
| *D. enghoffi* | 0.67997 | 0.00601 | 0.01530 | 0.03493 | 0.00033 | 0.06281 | 0.00000 | 0.00000 | 0.20065 |
| *D. esquivelli* | 0.09166 | 0.37993 | 0.02091 | 0.03292 | 0.07623 | 0.00206 | 0.05285 | 0.00124 | 0.32368 |
| *D. gaifa* | 0.23104 | 0.00710 | 0.00264 | 0.35837 | 0.01032 | 0.13678 | 0.00001 | 0.00000 | 0.25373 |
| *D. gollumi* | 0.16034 | 0.21028 | 0.61287 | 0.00000 | 0.00000 | 0.00000 | 0.00442 | 0.00025 | 0.01183 |
| *D. guayota* | 0.15824 | 0.00096 | 0.00064 | 0.45649 | 0.00327 | 0.27779 | 0.00000 | 0.00000 | 0.10261 |
| *D. hernandezi* | 0.00351 | 0.39558 | 0.02222 | 0.00001 | 0.00084 | 0.00000 | 0.50454 | 0.04387 | 0.00669 |
| *D. madai* | 0.38648 | 0.07878 | 0.03254 | 0.03284 | 0.00557 | 0.01287 | 0.00063 | 0.00001 | 0.45021 |
| *D. mahan* | 0.17166 | 0.01128 | 0.00238 | 0.42327 | 0.02309 | 0.10354 | 0.00004 | 0.00000 | 0.26470 |
| *D. minutisima* | 0.91765 | 0.00111 | 0.02607 | 0.00161 | 0.00000 | 0.02028 | 0.00000 | 0.00000 | 0.03328 |
| *D. ratonensis* | 0.18713 | 0.35679 | 0.05989 | 0.00796 | 0.01139 | 0.00130 | 0.02123 | 0.00050 | 0.35137 |
| *D. sibyllina* | 0.44086 | 0.01547 | 0.01081 | 0.10892 | 0.00361 | 0.06750 | 0.00002 | 0.00000 | 0.35279 |
| *D. unguimanis* | 0.00107 | 0.25398 | 0.00622 | 0.00001 | 0.00190 | 0.00000 | 0.62130 | 0.05477 | 0.00344 |
