## Supplementary material for "Tempo and mode of diversification of the red devil spiders (Araneae: Dysderidae) of the Canary Islands": Table S2

Table S2: Binari states cheliceral combinations that has shown significant differences between their diversification rates.

| A | B | C | D | E | F | G | I | λ0 | λ1 | p-value |
| --- | --- | --- | --- | --- | --- | --- | --- | --- | --- | --- |
| 0 | 1 | 1 | 0 | 0 | 1 | 0 | 0 | 0.01467329 | 0.049045711 | 0.00241025 |
| 0 | 1 | 0 | 0 | 1 | 1 | 0 | 0 | 0.010343693 | 0.049264661 | 0.00040656 |
| 0 | 1 | 1 | 0 | 1 | 1 | 0 | 0 | 0.010757809 | 0.046667713 | 0.00097059 |
| 0 | 1 | 0 | 0 | 0 | 0 | 1 | 0 | 0.016900501 | 0.044228815 | 0.01433236 |
| 0 | 1 | 1 | 0 | 0 | 0 | 1 | 0 | 0.017557185 | 0.042202741 | 0.02465853 |
| 0 | 0 | 0 | 0 | 1 | 0 | 1 | 0 | 0.012911226 | 0.056440032 | 0.00959059 |
| 0 | 1 | 0 | 0 | 1 | 0 | 1 | 0 | 0.012090296 | 0.042890574 | 0.00604434 |
| 0 | 1 | 1 | 0 | 1 | 0 | 1 | 0 | 0.011975284 | 0.041307962 | 0.01647841 |
| 0 | 1 | 0 | 0 | 0 | 1 | 1 | 0 | 0.018388807 | 0.042573488 | 0.01561464 |
| 0 | 1 | 0 | 0 | 1 | 1 | 1 | 0 | 0.013170213 | 0.04161216 | 0.01420913 |
| 0 | 0 | 1 | 0 | 0 | 0 | 0 | 1 | 0.016663142 | 0.068813008 | 0.00121277 |
| 0 | 1 | 0 | 0 | 0 | 1 | 0 | 1 | 0.013749689 | 0.05151426 | 0.00101071 |
| 0 | 0 | 1 | 0 | 0 | 1 | 0 | 1 | 0.014562464 | 0.06327448 | 0.0001861 |
| 0 | 1 | 1 | 0 | 0 | 1 | 0 | 1 | 0.01490424 | 0.048240716 | 0.00312824 |
| 0 | 0 | 0 | 0 | 0 | 0 | 1 | 1 | 0.014624123 | 0.054290233 | 0.00070946 |
| 0 | 1 | 0 | 0 | 0 | 0 | 1 | 1 | 0.017611685 | 0.043394353 | 0.02108051 |
| 0 | 0 | 1 | 0 | 0 | 0 | 1 | 1 | 0.015393235 | 0.050277293 | 0.001908 |
| 0 | 1 | 1 | 0 | 0 | 0 | 1 | 1 | 0.018166296 | 0.04154504 | 0.0322087 |
| 0 | 0 | 0 | 0 | 1 | 0 | 1 | 1 | 0.013261068 | 0.053205225 | 0.00502887 |
| 0 | 1 | 0 | 0 | 1 | 0 | 1 | 1 | 0.012606966 | 0.042255608 | 0.00883915 |
| 0 | 0 | 1 | 0 | 1 | 0 | 1 | 1 | 0.013490323 | 0.048113714 | 0.00500981 |
| 0 | 1 | 1 | 0 | 1 | 0 | 1 | 1 | 0.012471481 | 0.040800695 | 0.03907778 |
| 0 | 1 | 0 | 0 | 0 | 1 | 1 | 1 | 0.01904088 | 0.041851796 | 0.03364306 |
| 0 | 0 | 1 | 0 | 0 | 1 | 1 | 1 | 0.016528723 | 0.048041055 | 0.00492337 |
| 0 | 1 | 0 | 0 | 1 | 1 | 1 | 1 | 0.013699994 | 0.041096663 | 0.03531138 |
